# An ensemble deep learning framework to refine large deletions in linked-reads

**DOI:** 10.1101/2021.09.27.462057

**Authors:** Yunfei Hu, Sanidhya V Mangal, Lu Zhang, Xin Zhou

## Abstract

The detection of structural variants (SVs) remains challenging due to inconsistencies in detected breakpoints and biological complexity of some rearrangements. Linked-reads have demonstrated their superiority in diploid genome assembly and SV detection. Recently developed tools Aquila and Aquila_stLFR use a reference sequence and linked-reads to generate a high quality diploid genome assembly, using which they then detect and phase personal genetic variations. However, they both produce a substantial proportion of false positive deletion SV calls. To take full advantage of linked-reads, an effective downstream filtering and refinement framework is needed pressingly. In this work, we propose AquilaDeepFilter to filter large deletion SVs from Aquila and Aquila_stLFR. AquilaDeepFilter relies on a deep learning ensemble approach by integrating six state-of-the-art CNN backbones. The filtering of deletion SVs is formulated as a binary classification task on image data that are generated through the extraction of multiple alignment signals, including read depth, split reads and discordant read pairs. Three linked-reads libraries sequenced from the well-studied sample NA24385 and the gold standard of GiaB benchmark were used to perform thorough experiments on our proposed method. The results demonstrated that AquilaDeepFilter could increase the precision rate of Aquila while the recall rate of Aquila decreased only slightly, and the overall F1 improved by 20%. Furthermore, AquilaDeepFilter outperformed another deep learning based method for SV filtering, DeepSVFilter. Even though we designed AquilaDeepFilter for linked-reads, the framework could also be used to improve SV detection on short reads.

## I. Introduction

Structural variants (SVs) are large scale DNA alterations, including deletions, insertions, inversions and translocations covering at least 50 base pairs [1]. SVs could have a causal relationship with genetic diseases [2], as do small scale point mutations, single-nucleotide polymorphisms (SNPs) which are far more often characterized in genetic studies. SVs are less common than SNPs, but because they involve greater disruptions, they collectively amount for roughly as many bases in our genomes as SNPs and are therefore predicted to have a significant phenotypic impact [3]. Accurate detection of SVs could further explain the etiology of diseases and aid diagnosis. With current advancements in whole genome sequencing (WGS) [4] complemented by a boom in the compute power of GPUs and CPUs, the way has been paved for exploration of brand-new SV detection methods and optimization of prior practices.

SVs are difficult to detect by conventional sequencing methods. Existence of breakpoints may result in inaccurate alignment of reads. However, SVs could also generate many unique alignment signals that could be exploited to assist detection, including changes in read depth, discordant read pairs and split reads [5]. Many start-of-the-art SV callers design algorithms with the assistance of these divergent signals to detect SVs. There are over 40 SV callers and algorithms published since 2010 [6]. Methods for SV calling from WGS data mostly fall into two categories: alignment-based and assembly-based. Alignment-based approaches align reads into the reference genome and determine the location in the genome of the specific read’s best matches. Alignment-based methods can be applied to discover SNPs, small insertions and deletions (indels) and large SVs. The accuracy of SNP and small indel calling is near perfect in short reads WGS, whereas accuracy is much poorer for SVs [7]. Assembly-based methods assemble the full length of sequences (contigs) first, and then seek to detect variations by aligning to the reference. Alignment-based approaches generally enable the detection of SV at better resolution and higher sensitivity, by avoiding reference bias from alignment [8]. However, the computation cost is generally higher than alignment-based approach.

In recent years, deep learning methods have gained more popularity in various research fields including variant detection in WGS. DeepVariant started the trend of combining an image processing workflow with the small variant detection problem [9]. By using the Inception Convolutional Neural Network (CNN) model to classify the pileup aligned reads around the candidate variants as images, DeepVariant outperformed all other existing algorithms and tools of detecting SNPs, and small indels. DL-CNV [10] mostly focuses on the exon read depth signal, generating images and taking a similar strategy as DeepVariant to detect copy number variations (CNVs). CNNScoreVariants [11], a downstream filtering module of the GATK toolbox, encodes additional signals like mapping quality and read flags to enhance calling results and optimizes the CNN structures as well. Clairvoyante [12] preserves the read and reference information to form the training images and outperforms DeepVariant [9] on PacBio Single Molecule Sequencing data with much less trainable parameters. DeepSV-Filter [13] incorporates divergent signals from alignment reads as 3-channel images for large deletion and duplication filtering, and likewise, it utilizes CNN models to increase the calling precision of upstream tools like Lumpy, delly [14] and manta [15]. Convolutional neural network-based methods, equipped with supervised learning approaches inspired by computer vision tasks, have exhibited unique potential for SV filtering and refinement by sifting out false positive results of upstream detection algorithms so that the precision rate could increase.

Linked-reads is an advancing high throughput sequencing technology that has the potential to enhance long range information by assigning a unique barcode for reads that originate from the same long DNA molecule while maintaining the advantages of short-reads strategy as low cost and error rate [16]. Linked-reads sequencing has demonstrated its superiority in genome assembly, SV detection and phasing [17]. Specifically designed for 10x and single tube long fragment read (stLFR) linked-reads data recently, Aquila [18] and Aquila_stLFR [19] are reference-assisted local assembly approach to generate high-quality diploid assembly and enable the genome-wide discovery and phasing of all types of variants including SNPs, small indels and SVs. They are hybrid approaches that use both an alignment signal and an assembly strategy to achieve a promising high sensitivity for SV calls in linked-reads, however, their precision is somewhat limited due to the existence of large amount of false positives, especially for large deletion SVs. Thus, filtering out false positives from SV calls will significantly improve the performance of linked-reads in characterizing SVs.

In this study, we propose a framework for SV filtering and refinement in linked-reads, AquilaDeepFilter, which can be applied to linked-reads WGS data. Multiple alignment divergent signals and encoding methods are utilized to enhance image quality and to reduce computation cost. By using an ensemble strategy, several state-of-the-art CNN backbones are selected, with the further modification of adding a buffering convolution layer to them. These are then pre-trained to further enhance the accuracy of SV filtering. We have run several experiments on three different linked-reads libraries to refine large deletion SV calls of Aquila and Aquila_stLFR and the result of the preceding SV filtering method is shown for comparison. The results show that AquilaDeepFilter significantly reduces false positives while it only suffers a slight decrease in recall rate. The F1 scores are generally increased by 20% in all experiments compared to the SV calling result of Aquila. The comparison result shows that AquilaDeepFilter substantially outperforms DeepSVFilter on linked-reads data on precision rate, recall rate and f1 score. Even though AquilaDeepFilter is only applied to linked-reads in this paper, this ensemble framework will be flexible to be applied to short-read data as a filtering and refinement framework for upstream SV callers.

## II. Methods

In this section, the general pipeline and its principal parts are delineated first. We subsequently describe the CNN backbone structures and corresponding modifications that are applied in AquilaDeepFilter to allow various sequencing coverages and flanking region lengths around each SV. Finally, we present the ensemble strategy to further enhance the prediction robustness and accuracy of our method.

### A. The pipeline

Aquila uses a reference sequence and linked-reads to generate a high quality diploid genome assembly, from which it then comprehensively detects and phases personal genetic variation. Three linked-reads libraries of the well-characterized NA24385 sample are used through Aquila variants calling because this sample has the gold standard provided by the Genome in a Bottle (GiaB) project [20]. We further utilize Truvari, a commonly used open source tool for benchmarking [21], with the aforementioned gold standard to evaluate SV calls from Aquila, and generate true positives (TPs), false negatives (FNs) and false positives (FPs). The positive and negative samples for AquilaDeepFilter are constructed from the TPs, FNs and FPs, respectively.

AquilaDeepFilter is a majority voting framework that assembles multiple state-of-the-art CNN classification backbones with a carefully designed first layer to satisfy customized input size. Based on the previous idea of regarding sequences as image signals, the reads information that is stored in the BAM file is encoded as RGB images with a customized height, which is consistent with sequencing coverage. Based on the strategies from previous studies, we have all three alignment divergent signals extracted and used them to assist filtering of SVs. Taking large deletion as an example here, the distance within a discordant read-pair would need to be marked as larger than expected due to the existence of a deletion SV. If the distance is abnormally larger than a given threshold and the reads are within the region of candidate deletion SV, the generated image will record this signal by assigning 1 to the B channel of the corresponding pixel. We use 200bp left and right flanking region around breakpoints to define the target region for each candidate SV. The read depth signal is encoded by assigning 1 to the R channel if the reads are mapped to the reference within the candidate region. One read could be split if a SV breaks the continuity when mapping to reference genome and will be documented with a soft-clip tag in the BAM file. The split read signal is stored if a read is soft-clipped following the same method in G channel of the image. The pipeline is illustrated in Fig. 1, which consists of 4 sequential procedures for training and prediction.

**Fig. 1:**
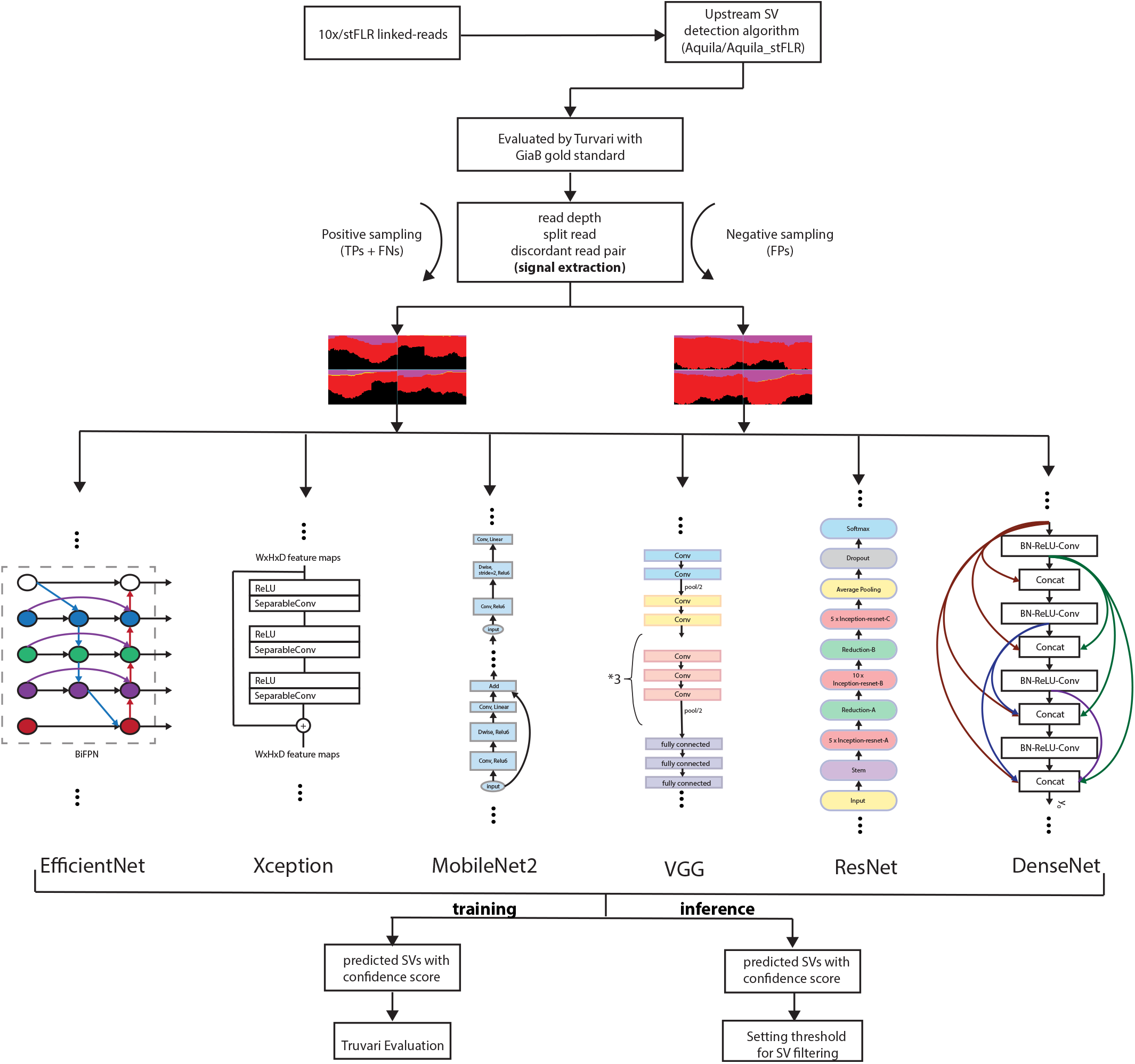
The pipeline of AquilaDeepFilter. There are six different SOTA CNN backbones modified and trained in this study, and a part of structure details are plotted as the representative of the whole model. The pipeline can be coarsely divided into 4 parts: raw calling result acquisition, image encoding, model training and evaluation/inference. Model training and evaluation shares the previous two phases and evaluation should be carried out after the models are well trained.

The initial step is the same for both training and inference, as during this phase we convert deletion SV calls of Aquila (or other SV detection algorithm) into BED files. The BED files serve as the position indicators that allow the workflow to extract the reads around those suspicious sites and thus extract several types of signals out of them. These BED files combined with the reference genome and aligned reads are exploited to generate the images, which will be passed to the ensemble to filter high confidence deletion SVs so that the algorithm’s precision rate could increase.

Once the candidate deletion SVs are identified, we proceed to image encoding, which is the second step in our pipeline. In this step we prepare the images that will be the input of our CNN models, details of which are discussed in subsequent sections. However, this step is slightly different for inference and training. In training mode, the images are split into two classes, positive and negative, to carry out supervised learning followed by data augmentation. Specifically, Truvari, a very common toolkit for benchmarking and annotating VCFs, is used here to acquire the labeled results of each called SV. The Truvari evaluation result of Aquila is divided into 4 parts, which are TPs, TNs, FPs and FNs, as mentioned. TPs and FNs are converted together to BED format as positive set while FPs are converted to negative set. In the training phase, the ground truth labels are assigned to each of the SV image but these labels are not generated in inference mode. Note that in the data augmentation step, based on the innate feature of signals extracted from DNA sequences, many possible image augmentation techniques cannot be applied since the generated images are less like natural images or pictures, which could tolerate rotation, color jittering, cropping or blurring. To the best of our knowledge, the only plausible augmentation technique here is flipping.

After the candidate deletion SVs are properly transformed to images, we proceed to the training/inference phase of pipeline. The CNN models are trained in a supervised manner and the prediction result of the model is stored in a BED file, which could be exploited to carry on extra downstream analysis or converted back to VCF format so the Truvari evaluation could be performed. The CNN model assigns a confidence score to each candidate deletion SV in BED file and a threshold could be set by the user to screen out the low confidence candidates. As is known, there is a trade-off existing in threshold setting since the higher threshold could increase the precision rate but decrease the recall rate.

Evaluation is the last part of the whole pipeline. It is used to estimate the generalization ability of the model on unseen data as it determines how well a model performs on new data. In our study, we use two different evaluation strategies in different phases. The cross-validation strategy is used in training phase where we split the training data into a ratio of 9:1 in training set and evaluation set respectively. And in the last part of training branch, we use Truvari to evaluate the performance of the trained models on a brand new dataset as a more generalized and reliable benchmark. For instance, we use three linked-reads libraries in this paper, 10x lib1 and 10x lib2 from 10x linked-reads, and stLFR lib from stLFR linked-reads. If the model is trained with 10x lib2, 10x lib1 and stLFR lib are recruited to perform the Truvari evaluation.

### B. CNN Model

The experiment was carried on six different state-of-the-art CNN models, XceptionNet [22], Resnet [23], EfficientNet [24], Densenet [25], MobileNetv2 [26], and VGG [27], pretrained on the ImageNet dataset [28]. To accelerate the training process, we adopted a transfer learning strategy. Transfer learning is a methodology where we reuse the previously trained model (on ImageNet) as a starting point for the new task. Transfer learning can be pretty handy by subsequently reducing the training time yet yielding better results on a small set of data points. All six CNN models mentioned except VGG and Densenet use residual connections. To improve the model performance, we inserted a convolution layer to have all shapes of customized input images prepared for the forward propagation of following CNN backbones. In addition to this, the output from these models was sent to a post processing block containing a fully-connected layer followed by a dropout layer and a final logits layer for classifying the image.

For better convergence of our models, we use a learning rate decaying strategy which divides the learning rate by 10 only when the loss stops decreasing for 3 continuous epochs and a early-stopping strategy that halts the training process after the learning rate decays for 5 times.

When training the models, the image dataset is split to 90% to 10% for training and validation. The Adam optimizer is used and 64 images are trained as a batch. The model converges after about 170 epochs of training and the loss and accuracy are documented at the end of each epoch. The learning rate is set to 0.001 initially. We selected SparseCategorialCrossEntropy with logits as the loss function for back propagation. The trained CNN models score each SV candidate during testing and further predicting. A customized threshold is set to filter out SV candidates.

### C. Customizing image size

Not much attention has been paid to the input size of previous image-based SV calling methods since this size is restricted or simply set to default by the pooling layer of CNN classification models. Most of these methods just assemble the SV signals to form images with default size, frequently, as a square of 224*224 pixels. The square mode could lead to loss of information if the height is less than the average read depth of our library. Also, if the width of image is set too large, most of the lower part of images is just white noise, which causes unnecessary waste of the GPU memory and computation time but contributes very little to performance. So in AquilaDeepFilter (Fig. 2), we proposed a new plug-in buffering layer that could be suitable for many kinds of customized input sizes with just limited modification to the existing CNN backbones. If the average coverage of a given library is about 100X and the flanking region length around breakpoints is set to 200 bps, SV images of 400*200 size will be generated by concatenating the left and right flanks of 2 breakpoints. The buffering convolution layer can be adjusted accordingly by simply setting the striding parameter to (2, 1) and kernel size to (3, 2) so that the model could work. This slight modification to the CNN backbone could increase the prediction accuracy while reducing unnecessary waste of computation.

**Fig. 2:**
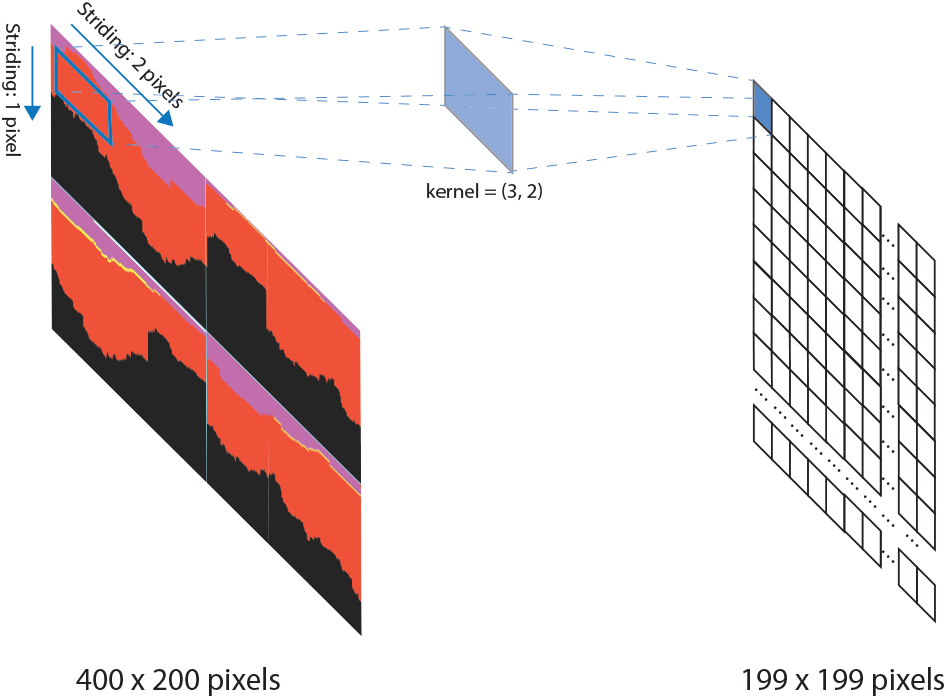
Buffering layer architecture. In order to make the input size consistent to the sequencing coverage and any customized change, the buffering layer serves as the “cushion” by setting proper kernel size and striding step to satisfy the needs. In our work, we choose a rectangular kernel of (3, 2) and striding step of (2, 1) to mitigate the loss of signal information as much as possible. The depth is not shown in this figure, but it could be adjusted flexibly by changing the number of kernels.

### D. Ensemble strategy

We noticed that during the experiment several CNN models had the consistent tendency to produce higher scores for those SVs that showed strong signals and were denoted as positive. However, for some of the candidates that were somewhat showing weaker signals or labeled as negative, the models rarely agreed with each other. This phenomenon led us to try the multi-model ensemble strategy, which is a widely used methodology where we combine multiple algorithms to obtain better predictive performance which could be hard to obtain from a single model. In this experiment we used the training accuracy acquired from the model training phase as the indicator for balancing the weights between several models, where predictions from all the models were computed and later a reduced mean operation was performed to obtain a consolidated result which is later used as a deciding factor whether the SV was positive or negative. This strategy here could greatly increase the precision rate of model while inhibits the loss of recall rate to some extent.

## III. Results

In order to illustrate the classification performance of DeepAquilaFilter, we have carried out a thorough investigation into the running time of model training and single instance inferring. The accuracy and robustness of AquilaDeepFilter, trained on two linked-reads datasets was tested with a 9:1 split ratio and carefully evaluated with Truvari by filtering out the deletion SV calling results of Aquila/Aquila_stLFR on other two linked-reads datasets, accordingly. This section will go over the experimental setup and the data we used, then the above results will be analyzed in sequential order.

### A. Experimental Setup

#### Linked-read libraries

In this study, three different linked-reads libraries sequenced from sample NA24385 are used to perform SV calling through Aquila/Aquila_stLFR and down-stream filtering for deletion SVs. The coverage of stLFR lib, 10x lib1 and lib2 are approximately 48X, 93X and 100X, respectively. Based on our previous studies [16], 10x libraries have better sequencing and assembly qualities than stLFR.

We used the Genome in a Bottle (GiaB) benchmark v0.6 as the gold standard in our work. This benchmark is specific to NA24385 and the hg19 reference sequence, with around 3600 positive deletion SVs ≥50 bp in high-confidence regions.

#### Dataset construction and training configuration

In this work, deletion SVs are detected by the state-of-the-art detection framework Aquila/Aquila_stLFR and are meticulously cleaned and selected according to the evaluation result of Truvari at beginning. The true positives and false negatives are divided into the positive set while the false positives are regarded as the negative set for AquilaDeepFilter. Table I shows the details of the composition of the training set. What follows next is the augmentation of the dataset. Unlike traditional types of image parsing tasks, the image that is generated in our study is less like natural images but more like integrated signals, so most of the augmentation techniques are not suitable and the only augmentation we performed is horizontal and vertical flipping. After the augmentation, the total number of all the training images had doubled.

**TABLE I:** Dataset statistics. The number of SV image with ground-truth label used for training.

| library | #SV images in training set |  |  |  |  |
| --- | --- | --- | --- | --- | --- |
|  | TP | FN | FP | positives | negatives |
| 10x lib1 | 3700 | 498 | 6870 | 4198 | 6870 |
| stLFR lib | 3277 | 921 | 8925 | 4198 | 8925 |
| 10x lib2 | 3349 | 322 | 6455 | 3671 | 6455 |

As for the validation set, since all three linked-reads libraries are from NA24385 and we can use the gold standard by GiaB to evaluate all of them, to generate TPs, FNs and FPs to create the positive and negative samples. The details of validation set is also illustrated in Table I.

Fig. 3 shows the basic settings of our experiments. We have carried three sets of experiments in total, while the first two experiments had two separate parts with different goals. As for the first two experiments, we train the AquilaDeepFilter on 90% and validate it on the rest 10% for curves plotting and hyperparameter adjustment. To achieve better performance on Truvari evaluation result, 100% data of each library is used for training and then the other two linked-reads datasets are exploited for evaluation.

**Fig. 3:**
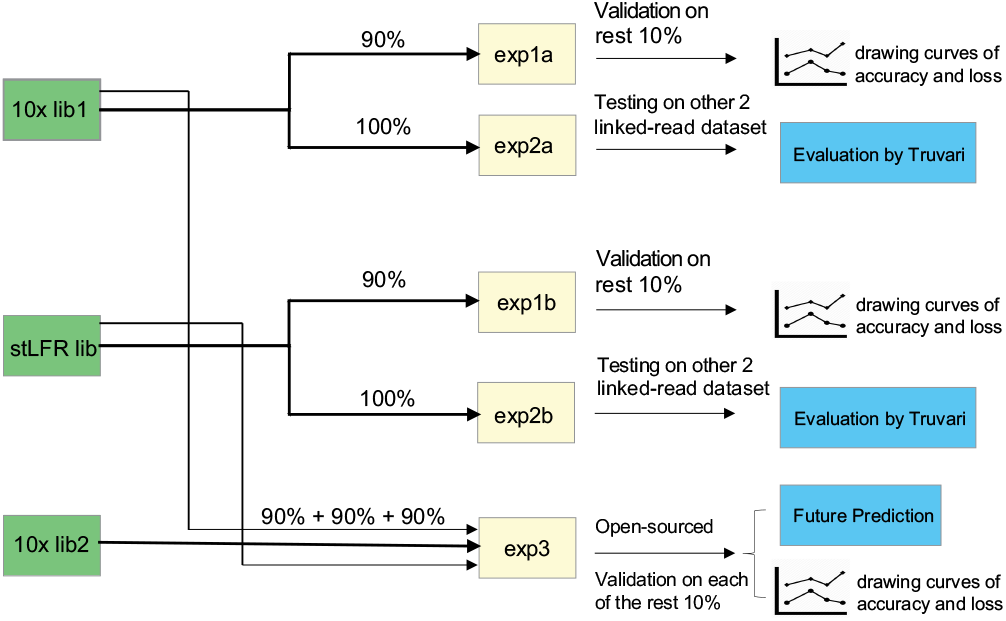
The logic flow chart of the main experiments. Since the 10x lib1 and lib2 are similar to some extent, we choose 10x lib1 and stLFR lib for model performance evaluation. In exp1a and exp1b, we train the AquilaDeepFilter on 90% of corresponding library, validate it on the rest 10% and plot the curves for accuracy and loss during training. In exp2a and exp2b, 100% of the lib data is used for Truvari evaluation. The weights of exp3 will be saved for future filtering of deletion SVs.

### B. Model Runtime Performance

Table II shows the training and inference time of CNN models of exp1a from Fig. 3. For all the experiments, we use a server with Intel(R) Xeon(R) CPU E5-2420 0 @ 1.90GHz, 24 cores and two high performance GPU RTX 2080 Ti. Approximately 177 seconds were needed when training each epoch of EfficientNet, which resulted to about 7 hours until convergence is reached. In the inference phase, GPU-enabled detection was not significantly fast while predicting single SV, however, filtering out all the SV candidates in a batch was more efficient. The rest of the models’ runtime data can be found in Table II. Although the ensemble strategy is generally time-consuming, it does not take too much time in this scenario. In total, the ensemble filtering result could be generated within 20 minutes on 2 separated linked-reads libraries with 6 different CNN models.

**TABLE II:** Model Runtime Performance. The ending epoch and time is documented if the early stopping strategy is triggered or the experiment reaches the maximum training epoch. The inference time is calculated by averaging the total runtime to the total number of instances. The training runtime data is from the experiment1a, and it would be longer in the experiment3 since it has essentially more training data.

| CNN Model | training |  | inference |
| --- | --- | --- | --- |
|  | convergence epochs | time | time/instance |
| ResNet | 169 | 748min | 7.91ms |
| EfficientNet | 146 | 432min | 3.63ms |
| VGG | 174 | 362min | 4.91ms |
| Xception | 173 | 856min | 8.78ms |
| MobileNetV2 | 180 | 483min | 4.87ms |
| DenseNet | 188 | 573min | 3.40ms |

### C. CNN model performance analysis

In order to demonstrate the performance of AquilaDeepFilter, the core components of this framework will be analyzed firstly. We employed 6 CNN backbone structures, which are DenseNet, ResNet, VGG, Xception, EfficientNet and MobileNetv2, and then modified them by adding a buffering layer to support a customized input image size. All the models were trained on the Linux server mentioned above.

We conducted the training of our models on the 10x lib2 and the stLFR lib, while the 10x lib1 was not used here because it had a similar sequencing quality and coverage as the 10x lib2. Both batches of models were trained with 90% of the training set and were validated with the rest 10% of the training set. Additionally, the model was trained on the mixed dataset of all three libraries, while 90% of each set was used for training. The evaluation metrics for performance evaluation are common and straightforward since the training task is merely a binary image classification problem. The accuracy is calculated as follows:

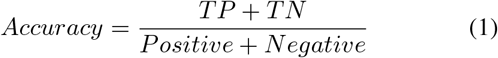

where *TP* (true positives) and *TN* (true negatives) are number of correctly classified instances, *Positive* and *Negative* are number of instances labelled positive and negative, respectively.

Fig. 4 shows the accuracy and loss curves for both the training and validation process of models trained on 10x lib2. Since the transfer learning strategy is taken, the initial loss is rather low and exhibits the potential to decrease until convergence. However, the accuracy and loss curves of the validation process oscillate from start to finish but also reveal the general tendency of stabilizing as epochs increase, gradually. The results confirm that the modified CNN models can efficiently fit the training data and show competitive classification ability on the validation set. The proposed method is assumed to be feasible for deletion SVs filtering in upstream calling result.

**Fig. 4:**
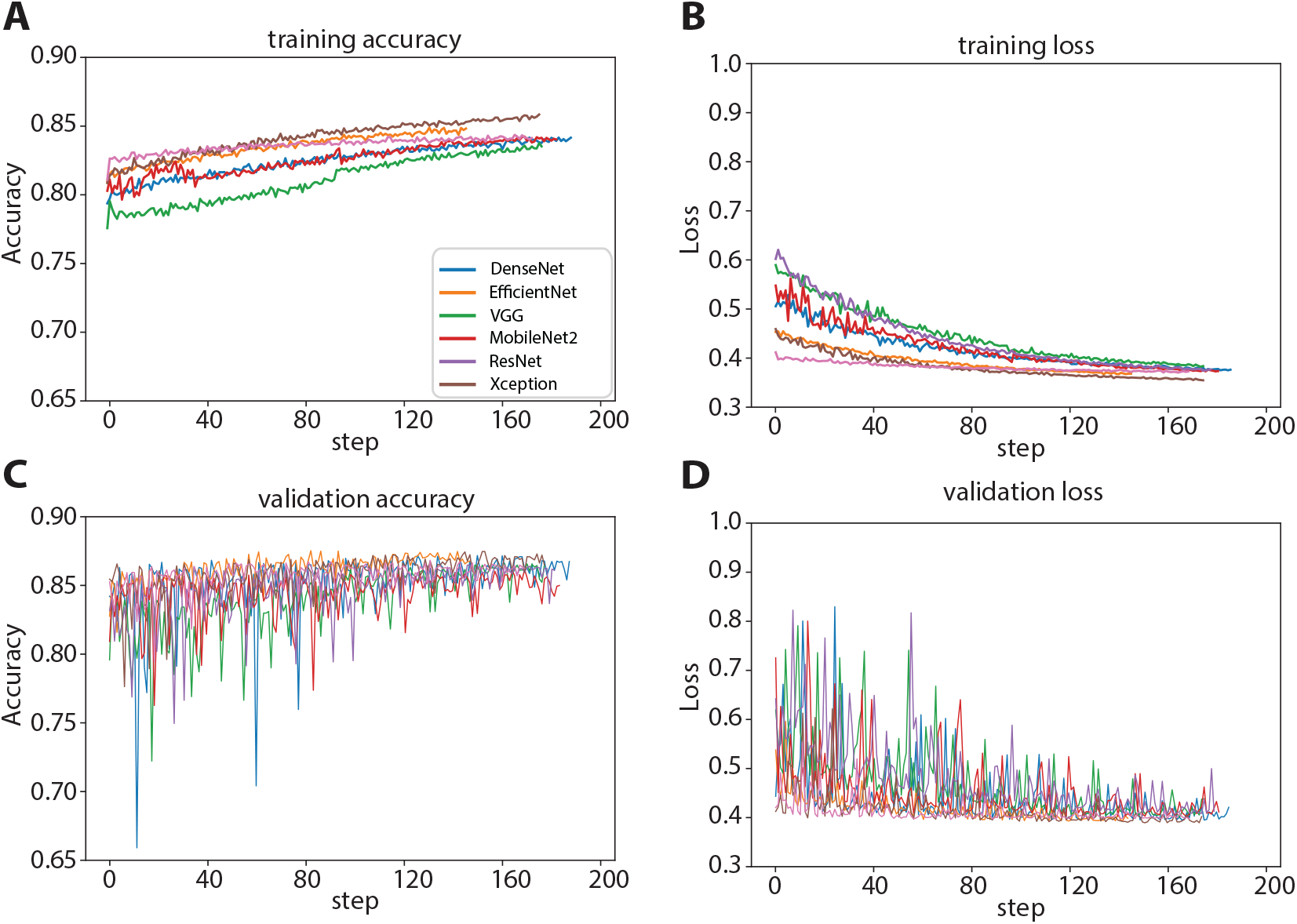
The training and loss curves of CNN models in AquilaDeepFilter trained on 10x lib2 data. A) and C) exhibits the accuracy of training process and validation after each training epoch. B) and D) shows the loss curves of training and validation, accordingly.

The remaining training and loss curves of models trained with all three linked-reads libraries and with the stLFR lib can be found in the Supplementary Fig. 1 and 2.

### D. AquilaDeepFilter performance analysis

In this part, we have performed thorough analysis on two trained models with the Truvari evaluation method. For simplicity, we only chose the corresponding experiment results of AquilaDeepFilter trained with the 10x lib2 data. The filtering experiment was carried on the remaining two linked-reads libraries other than the one that was already used for training. It is designed scrupulously for separating data for training with the data for Truvari evaluation, since we noticed that models trained and tested on the dataset with similar distribution could give better performance but often fail to work well on new data.

The filtering result of AquilaDeepFilter is shown in Table III while the separate filtering results of each CNN model can be found in the Supplementary Table 1 and 2. Based on the filtering results, the receiver operating characteristic (ROC) curves of all CNN models and AquilaDeepFilter are exhibited in Fig. 5. On the left side of Fig. 5, the ROC curves of filtering results on 10x lib1 data are plotted, while curves for stLFR lib are shown on the right. By comparing and analyzing the prediction scores of each candidate SV, we noticed that some candidates receive distinct scores from different models while most of the candidates shared similar scores. Thus, the ensemble strategy that aims to alleviate this issue proposed and the ROC curve of AquilaDeepFilter validates that ensemble strategy does receive higher AUC score and can classify more robustly. However, the ROC curves acquired from the models trained on stLFR lib are somewhat flat, comparably. This is a normal phenomenon due to the difference of two libraries’ sequencing quality. The lower sequencing quality and the error-prone characteristic of the data could disturb the generated images in a cascading manner and thus suppress the model performance.

**TABLE III:** AquilaDeepFilter evaluation result (trained on 10x lib2). The evaluation result on 10x lib1 is shown on the upper side of the table and result on stLFR lib is shown on the lower side.

| 10x lib1 | threshold | TP | FP | FN | precision | recall | f1 |
| --- | --- | --- | --- | --- | --- | --- | --- |
| Aquila | - | 3,700 | 6,870 | 498 | 0.35 | 0.8814 | 0.5011 |
| DeepSVFilter | 0.1 | 3,544 | 4,627 | 654 | 0.4337 | 0.8442 | 0.5730 |
|  | 0.2 | 3,373 | 3,048 | 825 | 0.5253 | 0.8035 | 0.6353 |
|  | 0.3 | 3,236 | 2,058 | 962 | 0.6113 | 0.7708 | 0.6818 |
|  | 0.4 | 3,064 | 1,408 | 1,134 | 0.6852 | 0.7299 | 0.7068 |
|  | 0.5 | 2,899 | 928 | 1,299 | 0.7575 | 0.6906 | 0.7225 |
|  | 0.6 | 2,707 | 594 | 1,491 | 0.8201 | 0.6448 | 0.7220 |
|  | 0.7 | 2,503 | 364 | 1,695 | 0.8730 | 0.5962 | 0.7086 |
| AquilaDeepFilter | 0.1 | 3,621 | 3,732 | 577 | 0.4925 | 0.8626 | 0.6270 |
|  | 0.2 | 3,461 | 1,977 | 737 | 0.6364 | 0.8244 | 0.7183 |
|  | 0.3 | 3,321 | 1,117 | 877 | 0.7483 | 0.7911 | 0.7691 |
|  | 0.4 | 3,165 | 601 | 1,033 | 0.8404 | 0.7539 | 0.7948 |
|  | 0.5 | 2,968 | 307 | 1,230 | 0.9063 | 0.7070 | 0.7943 |
|  | 0.6 | 2,829 | 129 | 1,369 | 0.9564 | 0.6739 | 0.7907 |
|  | 0.7 | 2,642 | 2 | 1,556 | 0.9992 | 0.6293 | 0.7723 |
| stLFR lib | threshold | TP | FP | FN | precision | recall | f1 |
| Aquila | - | 3,277 | 8,925 | 921 | 0.2686 | 0.7806 | 0.3996 |
| DeepSVFilter | 0.1 | 3,205 | 8,083 | 993 | 0.2839 | 0.7635 | 0.4139 |
|  | 0.2 | 3,116 | 6,839 | 1,082 | 0.3130 | 0.7423 | 0.4403 |
|  | 0.3 | 3,004 | 5,656 | 1,194 | 0.3469 | 0.7156 | 0.4673 |
|  | 0.4 | 2,882 | 4,488 | 1,316 | 0.3910 | 0.6865 | 0.4983 |
|  | 0.5 | 2,727 | 3,466 | 1,471 | 0.4403 | 0.6496 | 0.5249 |
|  | 0.6 | 2,583 | 2,503 | 1,615 | 0.5079 | 0.6153 | 0.5564 |
|  | 0.7 | 2,366 | 1,648 | 1,832 | 0.5894 | 0.5636 | 0.5762 |
| AquilaDeepFilter | 0.1 | 3,241 | 4,694 | 957 | 0.4084 | 0.7720 | 0.5342 |
|  | 0.2 | 3,082 | 3,058 | 1,116 | 0.5020 | 0.7342 | 0.5962 |
|  | 0.3 | 2,905 | 2,190 | 1,293 | 0.5702 | 0.6920 | 0.6252 |
|  | 0.4 | 2,793 | 1,611 | 1,405 | 0.6342 | 0.6653 | 0.6494 |
|  | 0.5 | 2,634 | 1,364 | 1,564 | 0.6588 | 0.6274 | 0.6428 |
|  | 0.6 | 2,436 | 1,144 | 1,762 | 0.6804 | 0.5803 | 0.6264 |
|  | 0.7 | 2,248 | 1,029 | 1,950 | 0.6860 | 0.5355 | 0.6015 |

**Fig. 5:**
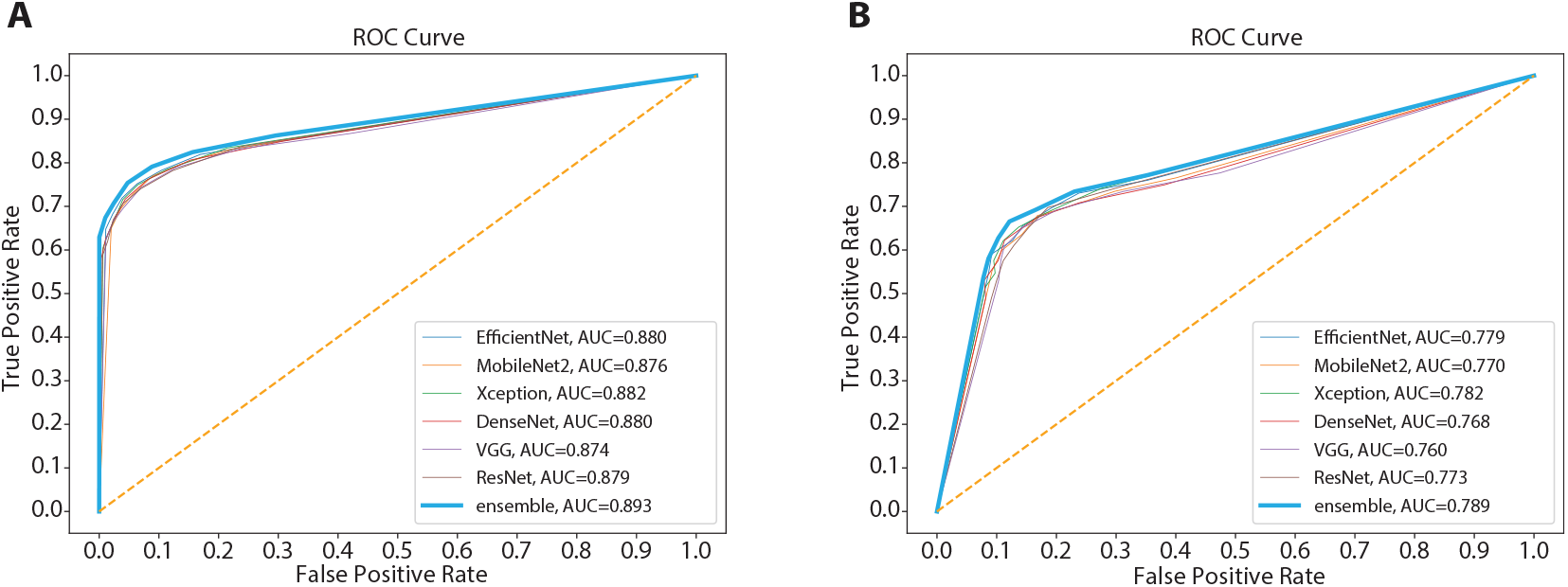
The ROC curves of AquilaDeepFilter and other CNN models trained on 10x lib2 linked-reads data. A) shows the ROC curve of Truvari evalution result on 10x lib1. B) shows the ROC curve of evaluation on stLFR lib.

Working as a downstream filter of deletion SVs called by Aquila/Aquila_stLFR, the fundamental issue that AquilaDeep-Filter should address is the low precision rate caused by genome-wide local-assembly strategy. In Table III, we used basic machine learning metrics to evaluate the deletion SV calling performance of Aquila/Aquila_stLFR and filtering result of AquilaDeepFilter. The precision, recall and f1 is calculated as follows:

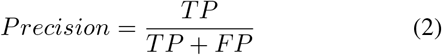

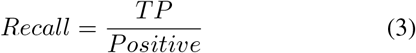

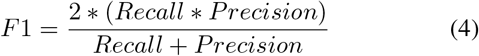

where *FP* (false positives) is the number of incorrectly classified instances.

The raw performance of Aquila reached 0.88 on 10x lib1 for deletion SV calls and Aquila_stLFR reached 0.78 on stLFR lib in terms of recall rate but showed a rather low precision rate. They could detect a high percentage of the potential deletion SVs but at the expense of mixing in large amount of false positives that are not easily validated.

With the help of AquilaDeepFilter, the precision rate of the filtered result immediately increased while the recall rate only decreased slightly. Although the libraries for training and Truvari evaluation differ from each other, AquilaDeepFilter does sufficiently learn the hidden features and patterns from the signals extracted from the BAM file, so that the F1 score can be greatly increased by filtering out large amount of negative candidates. In conclusion, AquilaDeepFilter has the capacity to serve as a downstream filtering and refinement framework for deletion SVs in linked-reads data. The overall performance of our framework could be further optimized if the training data and data for inference are of high quality.

We further did a comparison between DeepSVFilter and AquilaDeepFilter since both can work as a filtering step for upstream SV callers. The filtering result of DeepSVFilter is also shown in Table III. The same set of thresholds is used here for better comparison. We noticed that the CNN models used by DeepSVFilter were mostly light-weighted and had similar training performance. Given the fact that only the weights of MobileNet is open-sourced, we retrained this model with the same strategy on our linked-reads dataset as a benchmark. The result shows that AquilaDeepFilter outperforms DeepSVFilter in all three metrics.

## IV. Discussion and Conclusion

In this study, a novel deep learning-based framework, AquilaDeepFilter, is proposed for filtering deletion SVs in linked-reads WGS data. AquilaDeepFilter first extracts SV signals and integrates them as images. Secondly, after the training process, AquilaDeepFilter is applied to filter deletion SV calls detected by Aquila and Aquila_stLFR. Using three different linked-reads libraries, the performance of AquilaDeepFilter was thoroughly evaluated on the real WGS data of NA24385 with the gold standard provided by the GiaB project. AquilaDeepFilter achieved solid performance on reducing false positive rates while maintaining calling sensitivity as a downstream framework for linked-reads. Even though we run experiments for AquilaDeepFilter on deletion SV calls by Aquila/Aquila_stLFR, this ensemble model could be trained from SV calls by any upstream SV callers based on linked-reads or short-reads as a downstream framework for filtering and refinement.

We have adopted signal extraction methods of DeepSVFilter and Samplot-ML [29] which are thoughtfully designed and implemented. Novel contributions of our approach include the exploitation of more advanced CNN backbones and addition of a buffering layer that allows a customized input size rather than a fixed square input size. A further improvement over current methods involves performing the evaluation task both during the phase of neural networks training and testing with new linked-reads data from the same individual. This step demonstrated the generalization ability of our trained models on linked-reads data.

In spite of the aforementioned features, AquilaDeepFilter still has some limitations and the potential to evolve in the future. We have carried thorough experiments on deletion SV filtering since the methodology for extracting deletion signals is relatively mature in previous studies. There is still room for optimization for the insertions and other types of SVs, in terms of image processing from alignment signals. The filtering and detection of insertion, translocation and inversion SVs will be examined in future work.

## Supporting information

AquilaDeepFilter_SI

## Code Availability

We use the high-performance TensorFlow2 platform as the algorithm implementation framework. AquilaDeepFilter is freely available from the website at https://github.com/maiziezhoulab/AquilaDeepFilter.

## Funding

This work was supported by Vanderbilt University Development Funds (FF_300033). L.Z. is partially supported by Research Grant Council Early Career Scheme (HKBU 22201419).

## Acknowledgment

We wish to thank Yuanqi Xie for technical support on the manuscript.

## Notes

### Competing Interest Statement

The authors have declared no competing interest.

