## Supplementary material for "An ensemble deep learning framework to refine large deletions in linked-reads": AquilaDeepFilter_SI

**Hu et al.**

**Supplementary Information**

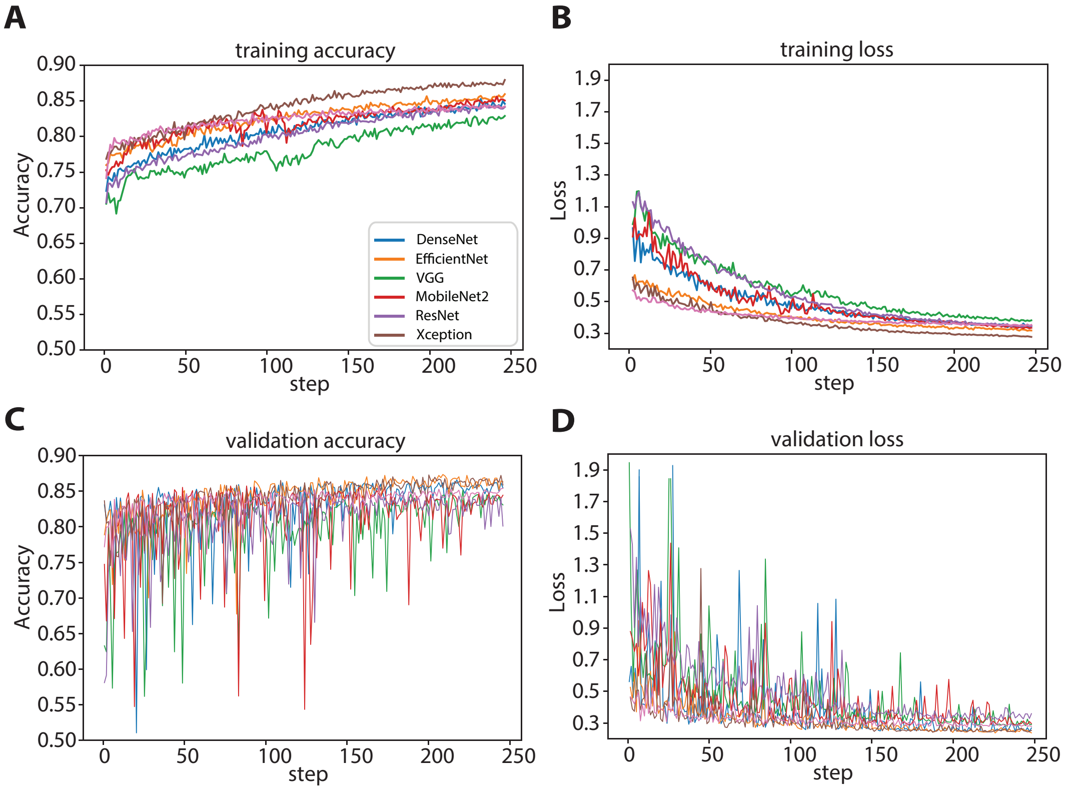


Supplementary Figure 1: The training and loss curves of AquilaDeepFilter and other models trained on mixed data

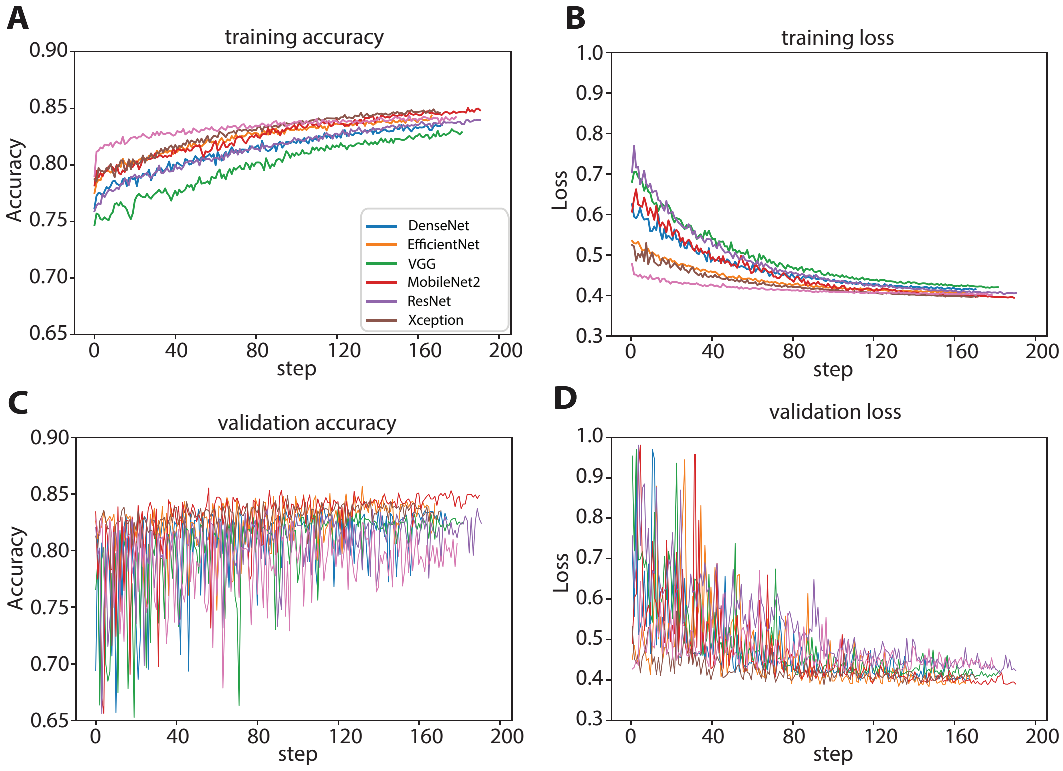


Supplementary Figure 2: The training and loss curves of AquilaDeepFilter and other models trained on stLFR lib data

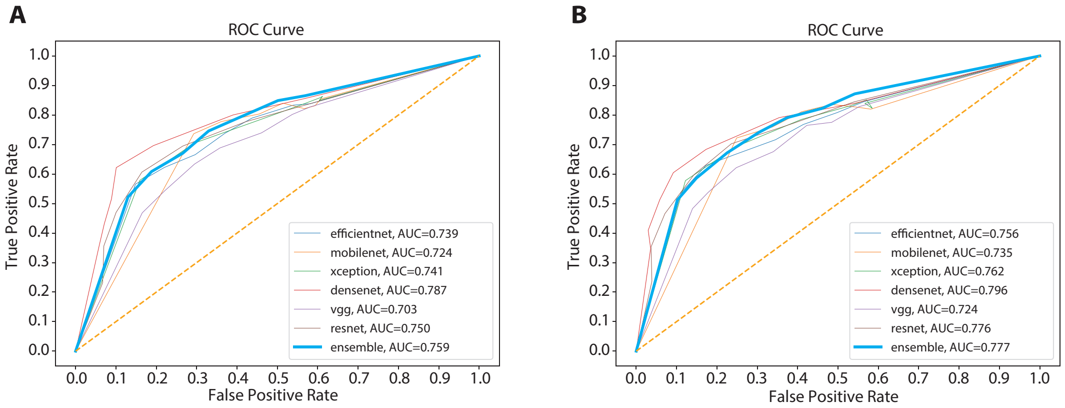


Supplementary Figure 3: The ROC curves of AquilaDeepFilter and other models trained on stLFR lib linked-reads data

Supplementary Table 1: 10x lib2 trained models evaluation results (on stLFR lib data)

| Model | threshold | TP | FP | FN | precision | recall | f1 |
|---|---|---|---|---|---|---|---|
| **Aquila** | - | 3,277 | 8,925 | 921 | 0.2686 | 0.7806 | 0.3996 |
| **EfficientNet** | 0.1 | 3,190 | 4,679 | 1,008 | 0.4054 | 0.7599 | 0.5287 |
| | 0.2 | 3,067 | 3,158 | 1,131 | 0.4927 | 0.7306 | 0.5885 |
| | 0.3 | 2,920 | 2,530 | 1,278 | 0.5358 | 0.6956 | 0.6053 |
| | 0.4 | 2,750 | 1,906 | 1,448 | 0.5906 | 0.6551 | 0.6212 |
| | 0.5 | 2,611 | 1,680 | 1,587 | 0.6085 | 0.6220 | 0.6151 |
| | 0.6 | 2,481 | 1,217 | 1,717 | 0.6709 | 0.5910 | 0.6284 |
| | 0.7 | 2,307 | 1,152 | 1,891 | 0.6670 | 0.5495 | 0.6026 |
| **MobileNet** | 0.1 | 3,214 | 5,322 | 984 | 0.3765 | 0.7656 | 0.5048 |
| | 0.2 | 3,084 | 3,956 | 1,114 | 0.4381 | 0.7346 | 0.5489 |
| | 0.3 | 2,928 | 2,879 | 1,270 | 0.5042 | 0.6975 | 0.5853 |
| | 0.4 | 2,850 | 2,234 | 1,348 | 0.5606 | 0.6789 | 0.6141 |
| | 0.5 | 2,669 | 1,890 | 1,529 | 0.5854 | 0.6358 | 0.6096 |
| | 0.6 | 2,516 | 1,444 | 1,682 | 0.6354 | 0.5993 | 0.6168 |
| | 0.7 | 2,318 | 1,224 | 1,880 | 0.6544 | 0.5522 | 0.5990 |
| **Xception** | 0.1 | 3,112 | 3,655 | 1,086 | 0.4599 | 0.7413 | 0.5676 |
| | 0.2 | 2,898 | 2,533 | 1,300 | 0.5336 | 0.6903 | 0.6019 |
| | 0.3 | 2,739 | 1,816 | 1,459 | 0.6013 | 0.6525 | 0.6258 |
| | 0.4 | 2,606 | 1,471 | 1,592 | 0.6392 | 0.6208 | 0.6298 |
| | 0.5 | 2,417 | 1,262 | 1,781 | 0.6570 | 0.5758 | 0.6137 |
| | 0.6 | 2,295 | 1,293 | 1,903 | 0.6396 | 0.5467 | 0.5895 |
| | 0.7 | 2,160 | 1,073 | 2,038 | 0.6681 | 0.5145 | 0.5813 |
| **DenseNet** | 0.1 | 3,147 | 5,066 | 1,051 | 0.3832 | 0.7496 | 0.5071 |
| | 0.2 | 2,973 | 3,163 | 1,225 | 0.4845 | 0.7082 | 0.5754 |
| | 0.3 | 2,831 | 2,221 | 1,367 | 0.5604 | 0.6744 | 0.6121 |
| | 0.4 | 2,613 | 1,504 | 1,585 | 0.6347 | 0.6224 | 0.6285 |
| | 0.5 | 2,419 | 1,364 | 1,779 | 0.6394 | 0.5762 | 0.6062 |
| | 0.6 | 2,225 | 1,051 | 1,973 | 0.6792 | 0.5300 | 0.5954 |
| | 0.7 | 2,060 | 1,084 | 2,138 | 0.6552 | 0.4907 | 0.5612 |
| **VGG** | 0.1 | 3,261 | 6,294 | 937 | 0.3413 | 0.7768 | 0.4742 |
| | 0.2 | 3,083 | 4,149 | 1,115 | 0.4263 | 0.7344 | 0.5395 |
| | 0.3 | 2,905 | 2,667 | 1,293 | 0.5214 | 0.6920 | 0.5947 |
| | 0.4 | 2,734 | 1,914 | 1,464 | 0.5882 | 0.6513 | 0.6181 |
| | 0.5 | 2,548 | 1,495 | 1,650 | 0.6302 | 0.6070 | 0.6184 |
| | 0.6 | 2,427 | 1,423 | 1,771 | 0.6304 | 0.5781 | 0.6031 |
| | 0.7 | 2,228 | 1,373 | 1,970 | 0.6187 | 0.5307 | 0.5714 |
| **ResNet** | 0.1 | 3,134 | 4,138 | 1,064 | 0.4310 | 0.7465 | 0.5465 |
| | 0.2 | 2,927 | 2,465 | 1,271 | 0.5428 | 0.6972 | 0.6104 |
| | 0.3 | 2,737 | 2,043 | 1,461 | 0.5726 | 0.6520 | 0.6097 |
| | 0.4 | 2,562 | 1,709 | 1,636 | 0.5999 | 0.6103 | 0.6050 |
| | 0.5 | 2,417 | 1,476 | 1,781 | 0.6209 | 0.5758 | 0.5975 |
| | 0.6 | 2,224 | 1,326 | 1,974 | 0.6265 | 0.5298 | 0.5741 |
| | 0.7 | 2,052 | 1,209 | 2,146 | 0.6293 | 0.4888 | 0.5502 |
| **Ensemble** | 0.1 | 3,241 | 4,694 | 957 | 0.4084 | 0.7720 | 0.5342 |
| | 0.2 | 3,082 | 3,058 | 1,116 | 0.5020 | 0.7342 | 0.5962 |
| | 0.3 | 2,905 | 2,190 | 1,293 | 0.5702 | 0.6920 | 0.6252 |
| | 0.4 | 2,793 | 1,611 | 1,405 | 0.6342 | 0.6653 | 0.6494 |
| | 0.5 | 2,634 | 1,364 | 1,564 | 0.6588 | 0.6274 | 0.6428 |
| | 0.6 | 2,436 | 1,144 | 1,762 | 0.6804 | 0.5803 | 0.6264 |
| | 0.7 | 2,248 | 1,029 | 1,950 | 0.6860 | 0.5355 | 0.6015 |

Supplementary Table 2: 10x lib2 trained models evaluation results (on 10x lib1 data)

| Model | threshold | TP | FP | FN | precision | recall | f1 |
|---|---|---|---|---|---|---|---|
| **Aquila** | - | 3,700 | 6,870 | 498 | 0.3500 | 0.8814 | 0.5011 |
| **EfficientNet** | 0.1 | 3,558 | 3,720 | 640 | 0.4889 | 0.8475 | 0.6201 |
| | 0.2 | 3,439 | 2,138 | 759 | 0.6166 | 0.8192 | 0.7036 |
| | 0.3 | 3,286 | 1,318 | 912 | 0.7137 | 0.7828 | 0.7466 |
| | 0.4 | 3,156 | 802 | 1,042 | 0.7974 | 0.7518 | 0.7739 |
| | 0.5 | 3,018 | 482 | 1,180 | 0.8623 | 0.7189 | 0.7841 |
| | 0.6 | 2,865 | 286 | 1,333 | 0.9092 | 0.6825 | 0.7797 |
| | 0.7 | 2,720 | 139 | 1,478 | 0.9514 | 0.6479 | 0.7709 |
| **MobileNet** | 0.1 | 3,614 | 4,392 | 584 | 0.4514 | 0.8609 | 0.5923 |
| | 0.2 | 3,480 | 2,704 | 718 | 0.5627 | 0.8290 | 0.6704 |
| | 0.3 | 3,345 | 1,721 | 853 | 0.6603 | 0.7968 | 0.7222 |
| | 0.4 | 3,232 | 1,130 | 966 | 0.7409 | 0.7699 | 0.7551 |
| | 0.5 | 3,056 | 712 | 1,142 | 0.8110 | 0.7280 | 0.7673 |
| | 0.6 | 2,917 | 458 | 1,281 | 0.8643 | 0.6949 | 0.7704 |
| | 0.7 | 2,727 | 254 | 1,471 | 0.9148 | 0.6496 | 0.7597 |
| **Xception** | 0.1 | 3,502 | 2,715 | 696 | 0.5633 | 0.8342 | 0.6725 |
| | 0.2 | 3,293 | 1,426 | 905 | 0.6978 | 0.7844 | 0.7386 |
| | 0.3 | 3,146 | 816 | 1,052 | 0.7940 | 0.7494 | 0.7711 |
| | 0.4 | 3,005 | 501 | 1,193 | 0.8571 | 0.7158 | 0.7801 |
| | 0.5 | 2,811 | 323 | 1,387 | 0.8969 | 0.6696 | 0.7668 |
| | 0.6 | 2,671 | 184 | 1,527 | 0.9356 | 0.6363 | 0.7574 |
| | 0.7 | 2,537 | 78 | 1,661 | 0.9702 | 0.6043 | 0.7448 |
| **DenseNet** | 0.1 | 3,565 | 3,869 | 633 | 0.4796 | 0.8492 | 0.6130 |
| | 0.2 | 3,385 | 1,932 | 813 | 0.6366 | 0.8063 | 0.7115 |
| | 0.3 | 3,198 | 1,027 | 1,000 | 0.7569 | 0.7618 | 0.7593 |
| | 0.4 | 3,008 | 551 | 1,190 | 0.8452 | 0.7165 | 0.7756 |
| | 0.5 | 2,810 | 313 | 1,388 | 0.8998 | 0.6694 | 0.7677 |
| | 0.6 | 2,622 | 135 | 1,576 | 0.9510 | 0.6246 | 0.7540 |
| | 0.7 | 2,438 | 46 | 1,760 | 0.9815 | 0.5808 | 0.7297 |
| **VGG** | 0.1 | 3,641 | 5,317 | 557 | 0.4065 | 0.8673 | 0.5535 |
| | 0.2 | 3,491 | 2,969 | 707 | 0.5404 | 0.8316 | 0.6551 |
| | 0.3 | 3,310 | 1,621 | 888 | 0.6713 | 0.7885 | 0.7252 |
| | 0.4 | 3,122 | 890 | 1,076 | 0.7782 | 0.7437 | 0.7605 |
| | 0.5 | 2,922 | 510 | 1,276 | 0.8514 | 0.6960 | 0.7659 |
| | 0.6 | 2,788 | 289 | 1,410 | 0.9061 | 0.6641 | 0.7665 |
| | 0.7 | 2,615 | 140 | 1,583 | 0.9492 | 0.6229 | 0.7522 |
| **ResNet** | 0.1 | 3,516 | 3,006 | 682 | 0.5391 | 0.8375 | 0.6560 |
| | 0.2 | 3,284 | 1,558 | 914 | 0.6782 | 0.7823 | 0.7265 |
| | 0.3 | 3,109 | 881 | 1,089 | 0.7792 | 0.7406 | 0.7594 |
| | 0.4 | 2,968 | 507 | 1,230 | 0.8541 | 0.7070 | 0.7736 |
| | 0.5 | 2,809 | 295 | 1,389 | 0.9050 | 0.6691 | 0.7694 |
| | 0.6 | 2,601 | 178 | 1,597 | 0.9359 | 0.6196 | 0.7456 |
| | 0.7 | 2,467 | 72 | 1,731 | 0.9716 | 0.5877 | 0.7324 |
| **Ensemble** | 0.1 | 3,621 | 3,732 | 577 | 0.4925 | 0.8626 | 0.6270 |
| | 0.2 | 3,461 | 1,977 | 737 | 0.6364 | 0.8244 | 0.7183 |
| | 0.3 | 3,321 | 1,117 | 877 | 0.7483 | 0.7911 | 0.7691 |
| | 0.4 | 3,165 | 601 | 1,033 | 0.8404 | 0.7539 | 0.7948 |
| | 0.5 | 2,968 | 307 | 1,230 | 0.9063 | 0.7070 | 0.7943 |
| | 0.6 | 2,829 | 129 | 1,369 | 0.9564 | 0.6739 | 0.7907 |
| | 0.7 | 2,642 | 2 | 1,556 | 0.9992 | 0.6293 | 0.7723 |
